# Isolation and Characterization of L-Glutaminase producing Bacteria

**DOI:** 10.1101/2020.10.28.358838

**Authors:** Rabia Saleem, Safia Ahmed

## Abstract

Being a significant protein L-glutaminases discovers potential applications in various divisions running from nourishment industry to restorative and cure. It is generally disseminated in microbes, actinomycetes, yeast and organisms. Glutaminase is the principal enzyme that changes glutamine to glutamate. The samples were gathered from soil of Taxila, Wah Cantt and Quetta, Pakistan for the isolation of glutaminase producing bacteria. After primary screening, subordinate screening was done which includes multiple testification such as purification, observation of morphological characters and biochemical testing of bacterial strains along with 16S rRNA sequence homology testing. Five bacterial strains were selected showing glutaminase positive test in screening, enzyme production via fermentation and enzymatic and protein assays. Taxonomical characterization of the isolates identified them as *Bacillus subtilis* U1, *Achromobacter xylosoxidans* G1, *Bacillus subtilis* Q2, *Stenotrophomonas maltophilia* U3 and *Alcaligenes faecalis* S3. The optimization of different effectors such as incubation time, inducers, carbon source, pH, and nitrogen source were also put under consideration. There was slight difference among incubation of bacterial culture, overall, 36 hours of incubation time was the best for glutaminase production by all the strains. Optimal pH was around 9 in *Achromobacter xylosoxidans* G1 and *Alcaligenes faecalis* S3, pH 6 in *Bacillus subtilis* U1, pH 8 in *Stenotrophomonas maltophilia* U3, pH 6-8 in *Bacillus subtilis* Q2. Best glutaminase production was obtained at 37°C by *Bacillus subtilis* U1and *Bacillus subtilis* Q2, 30°C for *Achromobacter xylosoxidans* G1, *Stenotrophomonas maltophilia* U3 and 25°C by *Alcaligenes faecalis* S3. The carbon sources put fluctuated effects on activity of enzyme in such a way that glucose was the best carbon source for *Bacillus subtilis* U1and *Bacillus subtilis* Q2, Sorbitol for *Achromobacter xylosoxidans* G1 and *Alcaligenes faecalis* S3 while xylose was the best for *Stenotrophomonas maltophilia* U3. Yeast extract and Trypton were among good nitrogen sources for *Achromobacter xylosoxidans* G1 and of *Bacillus subtilis* U1 respectively. Glutamine was the best inducer for *Bacillus subtilis* Q2, *Alcaligenes faecalis* S3 and *Stenotrophomonas maltophilia* U3, while lysine for *Achromobacter xylosoxidans* G1 and glycine act as good inducer in case of *Bacillus subtilis* U1. After implementation of optimal conditions microbial L-glutaminase production can be achieved and the bacterial isolates have a great potential for production of glutaminase enzyme and their applications.

## Introduction

Being the most important entity in multiple biological and non-biological processes, enzymes play crucial role in maintaining and sustaining industrial, commercial and economical products. They include all biochemical response and accelerate the rate of response without being expressing themselves in the last product [1].

L-Glutaminase is amidohydrolase enzyme belong to hydrolytic class which catalyzes the conversion of amino acid L-glutamine into L-glutamic acid, in the presence of water and releases ammonia. This enzyme plays an important role in nitrogen metabolism at cellular level. This enzyme is present in both microorganisms such as bacteria fungi and yeast as well as in macro-organisms, so this is ubiquitous in nature [2]. The probable sources may include animals, plants, bacteria, actinomycetes, yeast and fungi [3]. Numerous bacteria are involved in synthesis of extracellular and intracellular glutaminases such as *Bacillus sp*., *Pseudomonas*, *Actinobacterium sp*, and *E. coli.* [4].

The focal sources of fungal glutaminases are *Aspergillus sp*. and *Trichoderma sp*. [6]. The glutaminase producing actinomycetes includes different species [7]. Plant tissue are also used in the production but the evidences of extraction of plant’s glutaminases are not too much due to less feasible approaches [8]. Due to complex organization, animals are not well known in the field of enzymatic isolation from their tissues [10]. So, microbes are the main source of enzymes. The approaches towards enzymatic production involved fermentation techniques, mainly of two types i.e. solid-state fermentation (SSF) and submerged fermentation (SmF) [11, 12].

Catalysts represent 80% of the complete mechanical market. Requests for chemical utilized in ventures are expanding step by step so as to improve the procedures [13].

L-Glutaminase is considered to be the most important enzyme in food industries, for enhancing the taste and aroma of fermented food [14, 15]. L-Glutaminsae is a potential anticancer enzyme, flavor enhancer, an antioxidant, and utilized as biosensor for deciding glutamine and glutamate [16]. Due to importance of this enzyme the present research is carried out to isolate new bacterial strains for glutaminase production.

## Materials and methods

### Sample collection

For isolation of microorganism soil samples were collected from Taxila, Wah Cantt and Quetta, Pakistan, kept in aseptic conditions in the lab and stored at −4°C. One bacteria strain was also isolated from old hydrolyzed glutamine sample.

### Primary screening

#### Isolation and screening of microorganism

For the isolation of bacteria from soil samples rapid plate method technique were used. The plate media were prepared whose composition was NaCl 0.5, KCl 0.5, MgSO_4._7H_2_O. 0.5, KH_2_PO_4_ 1, FeSO_4_.7H_2_O 0.1, ZnSO_4_ 0.1gm/L, L-Glutamine 0.5 as Nitrogen source and phenol red (0.002 g/L) for indicator of glutaminase activity by bacteria. After sterilization of media soil samples were sprinkled on medium and plates were incubating at 37°C in a thermal incubator. Bacteria which showed change in color around the colony due to basic pH were considered as glutamine producers and further proceeded.

#### Production of Glutaminase by fermentation method

Five bacterial isolates were selected and used for production of L-Glutaminase in fermentation media. The bacterial strains were inoculated in 500 ml Erlenmeyer flasks in screening media containing g/L Glucose, 10.0; Glutamine 5.0; Na_2_HPO_4_.2H_2_O, 6.0; KH_2_PO_4_, 3.0; MgSO_4_, 0.49; CaCL_2_, 0.002 and incubated in a shaking incubator for 96-120 hours at 25-37°C for bacteria at 120 rpm. The sample was collected at regular interval of 24 hours and centrifuged at 6000 rpm for 15 min at 4°C. The supernatant containing enzyme was collected and stored at 4°C.

### Secondary screening

#### Purification of bacterial strains

Isolates of bacteria which showed best activity in rapid plate assay were chosen and further purified by streak plate methods.

#### Extracellular enzyme activity by zone of hydrolysis

Purified bacterial strains were checked for extracellular enzyme activity by zones of hydrolysis. Very few cells of bacteria were inoculated on mid of screening media plates. Change in color was detected after 24 hrs.

#### Extracellular enzyme activity by well diffusion method

Bacterial isolates cultivated on production media and cell free broth was inoculated in wells (100 μl) in the screening media plates and next day the color change of the medium was observed and zone of hydrolysis was recorded.

#### Enzyme assay

The reaction mixture of enzyme assay contains 0.5 ml of Tris. HCL buffer of pH 8.0, 0.5 ml of 100 mM L-Glutamine; a substrate of L-Glutaminase, 0.5 ml of crude enzyme and 0.5 ml of distilled water. The reaction mixture was then incubated in a water bath at 37+2°C for 30 min. After incubation 0.5 ml of Trichloroacetic acid (TCA) was added for termination of reaction. In 3.7 ml of distilled water in a separate test tube 0.1ml of reaction mixture was added and finally 0.2 ml of Nessler reagent was added for estimation of nitrogen present in a sample [17]. The absorbance was measured at 450 nm and compared with standard curve to calculate Unit of enzyme.

#### Protein Estimation

Presence of extracellular protein in the crude enzyme was estimated by lowery method [18]. Bovine serum albumin (BSA) was used as a standard.

### Bacterial Identification

For identification of bacterial strains gram staining, morphological characters and biochemical tests were performed.

#### Gram staining

The reagents used in gram staining was crystal violet, gram iodine, ethanol and safranin. The reaction time for all of these reagents were 1 min except ethanol which is kept for 5 seconds on the slide. The color of the strains was observed along with other features under the microscope.

#### Morphological characterization

The bacterial strains were morphologically characterized on nutrient agar plates. The strains were streaked and incubated overnight and colony morphology was assessed in terms of size, pigmentation, shape, margin, elevation and opacity.

#### Biochemical test

Different biochemical tests were performed for identification taxonomy of bacterial strains. These biochemical tests include indole production, urease, catalase, oxidase, starch hydrolysis, carbohydrates fermentation (glucose, sucrose, lactose, xylose, mannose, and maltose), mannitol salt agar, and citrate utilization test using standard protocol [19].

### Molecular characterization

#### DNA isolation

For the extraction of DNA, a modified phenol chloroform DNA extraction method was performed. DNA was re-suspended in 30 μl of TE buffer in the tube and mixed the DNA pallet through pipetting and stored at 4°C. Isolated DNA was checked by gel electrophoresis using 0.8 % agarose. The DNA purity was observed by using spectrophotometer with an absorbance in a ratio of A_260/280_.

#### Amplification of 16S rRNA gene

Genomic DNA extracted from different bacteria was amplified by using PCR (Labnet international, Model: MultiGene OptiMax, USA). Reaction cycles were optimized by checking the PCR cycle times, melting and annealing temperature using universal primer 27F and 1492 R. The PCR reaction was carried out in a final volume of 25μl containing 2.5μl 1x PCR buffer, 5 μl of dNTPs, 3 μl of 1mM MgSO₂, 0.75 μl (IU) Taq DNA Polymerase, 1 μl of 10 μM Forward primer, 1μl of 10 μM Reverse Primer, 2 μl of bacterial Template DNA and 10.75 μl double distilled H₂ O. Amplification of DNA was carried out in 30 cycles for 1 hour and 40 minutes. For which each cycle run 1^st^ 10 second at 98°C then next 45 seconds at 94°C, 50 seconds at 52°C and then cycle was run at 72°C for 1 minutes and finished with 10 minutes at 72°C for final extension. The PCR amplified products were confirmed with the help of agarose gel electrophoresis.

#### Sequencing of 16S rRNA gene

The sequencing 16S rDNA amplified products of all the samples of bacteria conducted commercially. The sequences were blast with the help of BLASTN on the site of National Center for Biotechnology Information (NCBI). The sequences were used to make phylogenetic tree of all the bacterial strains was made by the aid of maximum likelihood method which cleared about evolutionary basis of strains.

#### Optimization of culture conditions for L-Glutaminase production

Optimization of multiple conditions leading to production of glutaminase was done by estimating optimal conditions such as effect of incubation time, pH, carbon source, nitrogen source and inducers.

#### Effect of incubation time

Glutaminase production by bacterial isolates was done at 30°C in fermentation media and activity was measured after specific intervals (0, 1, 2, 3, 4 and 5 days).

#### Effect of pH

The condition of pH was optimized for the production of glutaminase by the selected bacterial isolate by running fermentation at different pH (6-9).

#### Effect of Temperature

Selected bacterial strains were optimized for the effect of temperature at 25°C, 30°C and 37°C.

#### Effect of carbon source

All the strains were checked for their enzyme producing ability at different carbon source addition such as glucose, sucrose, lactose, maltose, xylose and sorbitol (1 %) in the production media.

#### Effect of nitrogen source

The effect of 0.1% nitrogen source was measured. The potent nitrogen sources tested were Trypton, Yeast extract, Ammonium chloride and Sodium nitrate.

#### Effect of inducers

The glutaminase producing strains were checked for measurement of effects of different inducers like Glycine, Glutamine and Lysine on glutaminase production and glutaminase activity was measured after every 24 hrs of incubation.

## Results

The present study showed the isolation and characterization of bacterial isolates with the capability of producing L-glutaminase enzyme.

### Primary Screening

The primary screening is basically the rapid plate assay technique which utilize phenol red as pH indicator. Soil samples were inoculated in the media and colonies displaying pink color formation around the colonies were picked and purified and further again tested for enzyme production on individual strains. Twenty bacterial colonies gave positive result in purified form changing the color of media from yellow to pink (Fig 1). Five isolated Q2, U3, U1, S3 and G1 were selected for further analysis. After the plate rapid assay, the strains were also tested in liquid media exhibiting same results.

**Fig 1.**
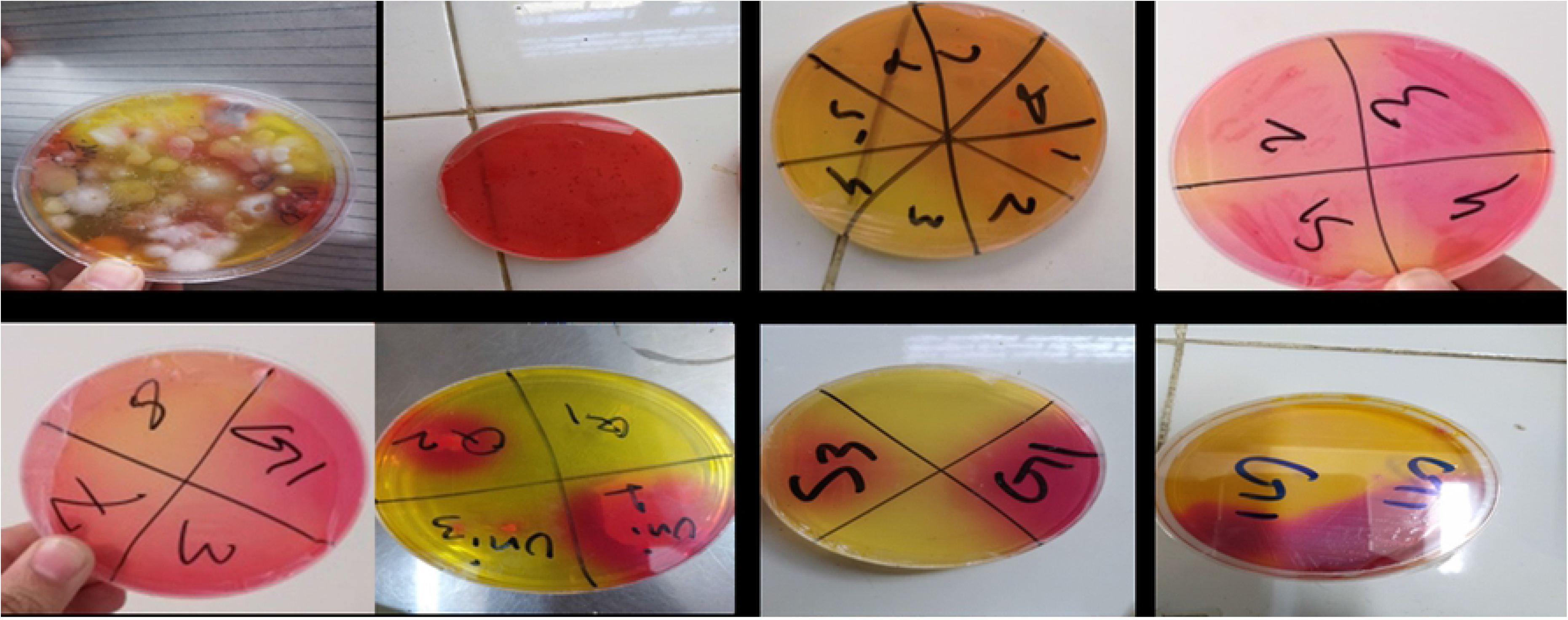
Primary Screening of bacteria in glutamine salt medium.

### Secondary screening

The secondary screening was done in media by point inoculation of bacterial strain in agar media and glutaminase production was indicated as color change in agar plates. The zone of hydrolysis was measured in millimeter, where glutaminase enzyme hydrolyze the substrate glutamine in a medium. The results showed that all the bacterial strain produce glutaminase giving zone of hydrolysis of 14-32 mm (Fig 2, Table 1). Further all the selected bacterial isolates were grown in liquid media (Fig 3) and cell free broth was added in the well for measuring activity of glutaminase in terms of dimeter of zone of hydrolysis in screening media. The results indicated that maximum diameter of zone of hydrolysis formed by strain *Stenotrophomonas maltophilia* U3 33 mm, *Achromobacter xylosoxidans* G1 was 26 mm, *Bacillus subtilis* U1 was 14 mm, Alcaligenes *faecalis* S3 was 12mm and *Bacillus subtilis* Q2 was 20 mm. While the activity of extracellular enzyme from other strains were less. The maximum zone of hydrolysis formed by *Bacillus subtilis* U1 was 34 mm in its diameter while strain *Alcaligenes faecalis* S3, *Achromobacter xylosoxidans* G1, *Stenotrophomonas maltophilia* U3 and *Bacillus subtilis* Q2 showed zone of hydrolyses of 30, 30, 23 and 16mm respectively (Fig 4, Table 1).

**Table 1.**
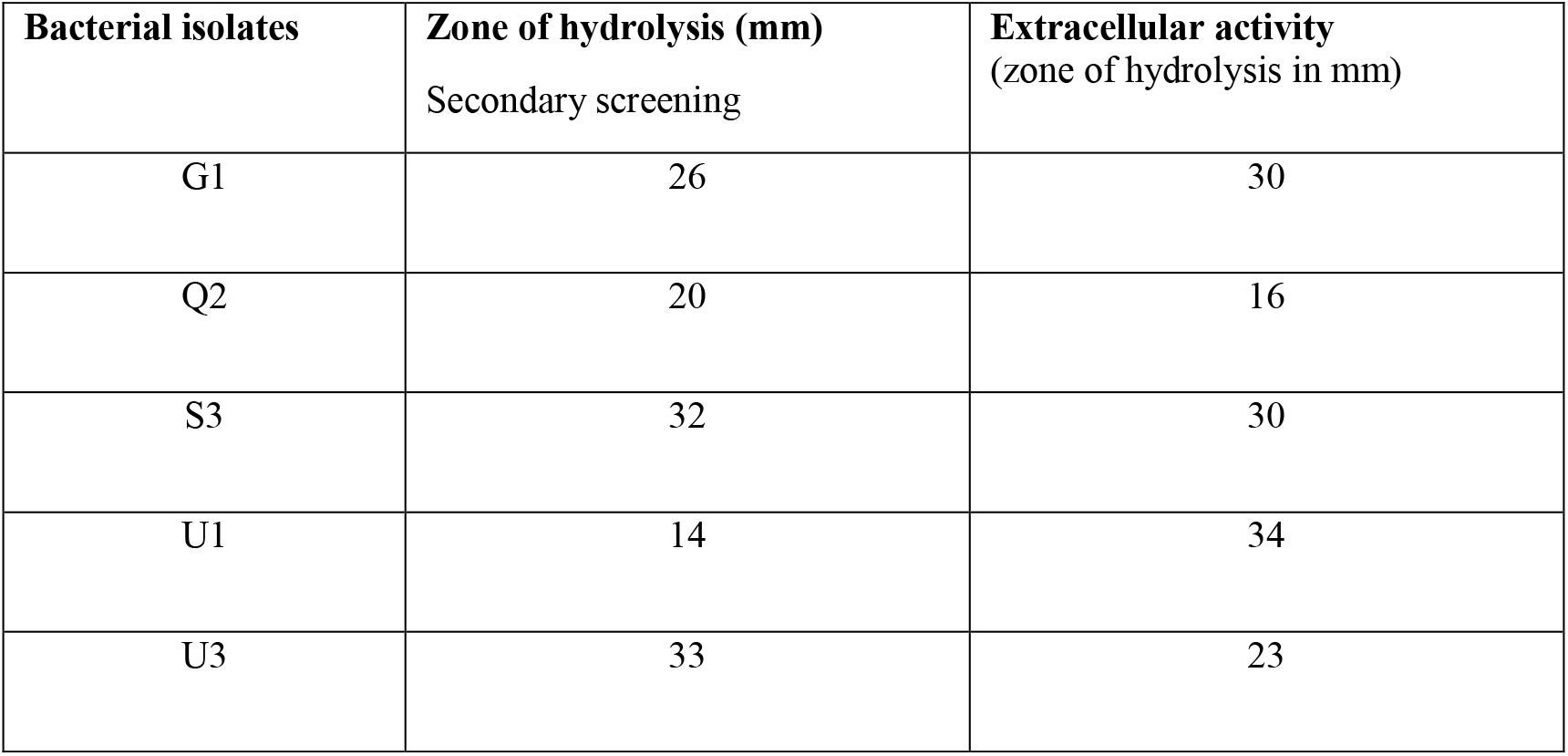
Screening of bacterial isolates for glutaminase production by Zone of hydrolysis.

| Bacterial isolates | Zone of hydrolysis (mm)<br>Secondary screening | Extracellular activity<br>(zone of hydrolysis in mm) |
| --- | --- | --- |
| G1 | 26 | 30 |
| Q2 | 20 | 16 |
| S3 | 32 | 30 |
| U1 | 14 | 34 |
| U3 | 33 | 23 |

**Fig 2.**
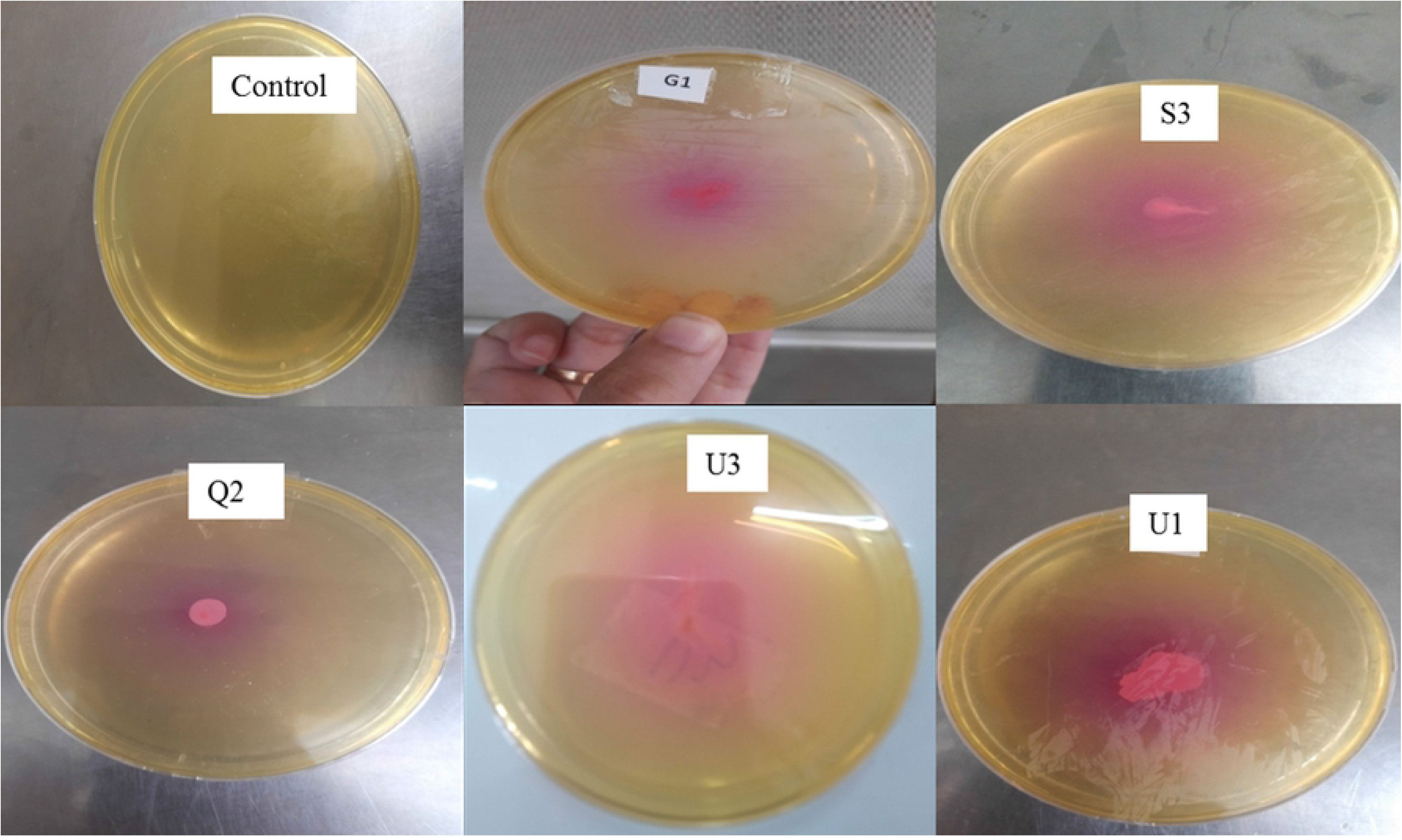
Zone of hydrolysis measured for L-glutaminase production by the bacterial isolates in screening media containing glutamine indicated as pink color.

**Fig 3.**
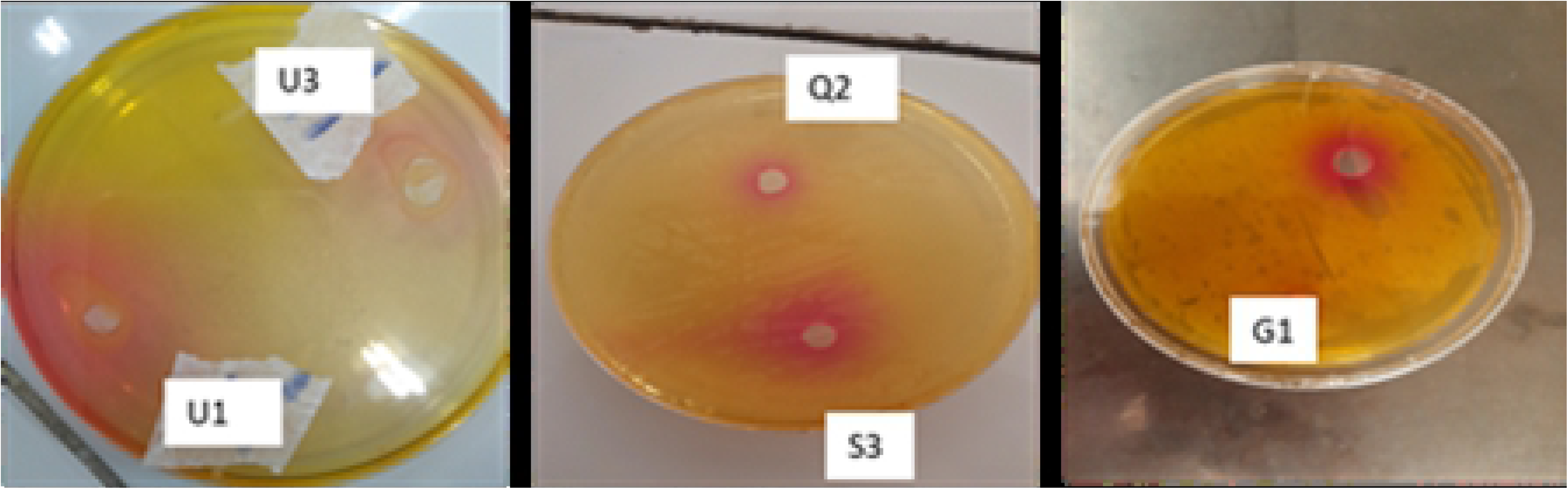
Extracellular enzyme screening on Glutamine Salt broth.

**Fig 4.**
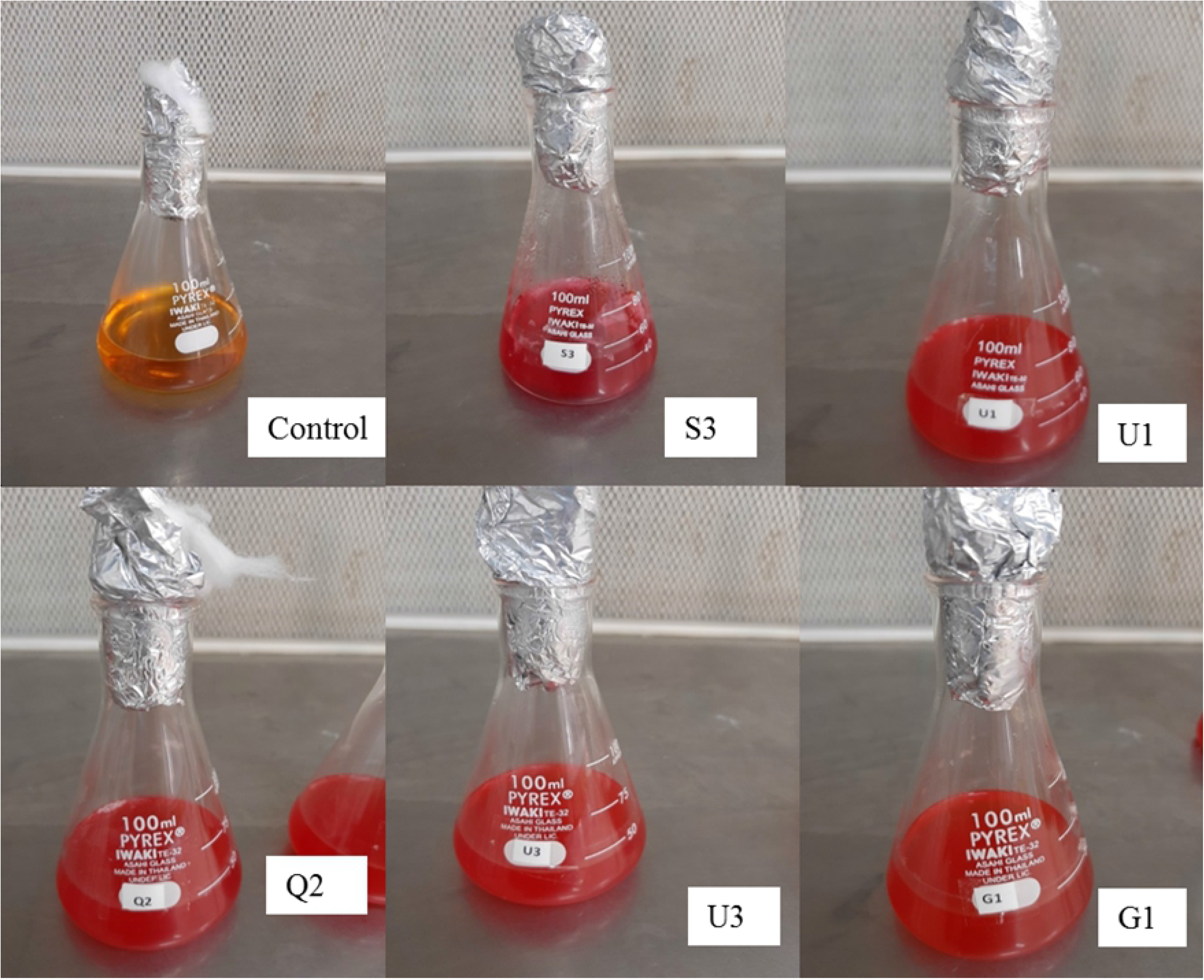
Extracellular enzyme activity by bacterial isolates in screening media using cell free broth in wells.

#### Taxonomic characterization of bacterial isolates

##### Cultural characteristics of bacterial isolates

The features like size, shape, margins, elevations, consistency, opacity and pigmentations showed slight differences in all the strains. Strain S3 showed small, G1 and U3 were of medium and U1 was having large sized colonies. As the shape of colonies is concerned, colonies of G1 strains were rhizoid in shape while circular shape exhibited by S3 and Q2 strains and U1 and U3 showed filamentous shape. All the strains were having dome shaped colonies except U1 which exhibited flat elevation. Colonies of all the strains were opaque and moist and showed no pigmentation (Table 2).

**Table 2.** Cultural characteristics of Bacterial isolates.

| Strains | Cultural characteristics |  |  |  |  |  |  |
| --- | --- | --- | --- | --- | --- | --- | --- |
|  | Size | Shape | Margins | Elevations | Consistency | Opacity | Pigmentation |
| **G1** | Medium | Rhizoids | Lobated | Dome | Moist | Opaque | Nil |
| **S3** | Small | Circular | Entire | Dome | Moist | Opaque | Nil |
| **Q2** | Large | Circular | Entire | Dome | Moist | Opaque | Nil |
| **U1** | Large | Filamentous | Filiform | Flat | Moist | Opaque | Nil |
| U3 | Medium | Filamentous | Filiform | Dome | Moist | Opaque | Nil |

#### Microscopic characters of the isolates

The microscopic observations such as shape, size and arrangement of cells was of different phases and features and the gram staining of strains revealed that all the strains were gram negative except U1 strains. The shape of the strain G1 was cocci while all others were rods (Table 3).

**Table 3.** Microscopic characters of bacterial isolates and Gram staining.

| Strains | Microscopic characters |  |  |  |
| --- | --- | --- | --- | --- |
|  | Shape | Size | Arrangement | Gram reaction |
| G1 | Cocci | Small | Single | Negative |
| S3 | Rod | Medium | Single | Negative |
| Q2 | Rod | Rod | Single | Negative |
| U1 | Rod | Rod | Single | Positive |
| U3 | Rod | Rod | Single | Negative |

### Biochemical testing

The biochemical tests were random except that of the strain U3 which was negative for all of the test and oxidase was positive for all of the test except U3. Strains G1, S3 and G2 were catalase positive and citrate utilization positive while Mannitol salt agar test was positive for G1, G2 and U1. Indole production was positive for G1 and Q2 and glucose fermentation was positive for S3, G2 and U1 while only strain G1 and S3 were urease positive (Table 4).

**Table 4.** Biochemical testing of bacterial isolates.

| Bacterial Isolates | Biochemical tests |  |  |  |  |  |  |
| --- | --- | --- | --- | --- | --- | --- | --- |
|  | Oxidase | Catalase | Mannitol | Citrate | Indole | Glucose | Urease |

|  |  |  | Salt agar<br>test |  | Production | fermentation |  |
| --- | --- | --- | --- | --- | --- | --- | --- |
| **G1** | + | + | + | + | + | – | + |
| **S3** | + | + | – | + | – | + | + |
| **Q2** | + | + | + | + | + | + | – |
| **U1** | + | – | + | – | – | + | – |
| **U3** | – | – | – | – | – | – | – |

### Molecular characterization

The agarose gel electrophoresis showed amplified sequences of 16S rRNA gene in bacterial isolates. The amplified product was of 1500 bp (Figure 5). The sequences were blast in NCBI database and on the basis of the homology to the sequences (> 90%) with other bacterial strains in the database the bacterial isolates were identified as *Bacillus subtilis* U1, *Achromobacter xylosoxidans* G1, *Bacillus subtilis* Q2, *Stenotrophomonas maltophilia* U3 (Table 5).

**Table 5.** 16S rDNA sequence homology and identification.

| **Strains** | **Size (bp)** | **Sequence homology with** | **Percentage homology** | **Accession number** | **Identified as** |
| --- | --- | --- | --- | --- | --- |
| U1 | 1467 | *Bacillus subtilis* strain B92 16S ribosomal RNA gene, partial sequence. | 98.32% | KT719942.1 | *Bacillus subtilis* U1 |
| S3 | 1369 | *Alcaligenes faecalis* strain SDU20 16S ribosomal RNA gene, partial sequence | 97.87% | MN229469.1 | *Alcaligenes faecalis* S3 |
| G1 | 805 | *Achromobacter xylosoxidans* strain BPS6 16S ribosomal RNA gene, partial sequence | 92.35% | MK786696.1 | *Achromobacter xylosoxidans* G1 |
| Q2 | 1200 | *Bacillus subtilis* strain B18 16S ribosomal RNA gene, partial sequence. | 97.87% | MK229037.1 | *Bacillus subtilis* Q2 |
| U3 | 1404 | *Stenotrophomonas maltophilia* strain NA156 16S ribosomal RNA gene partial sequence. | 96.56% | KT005287.1 | *Stenotrophomonas maltophilia* U3 |

**Fig 5.**
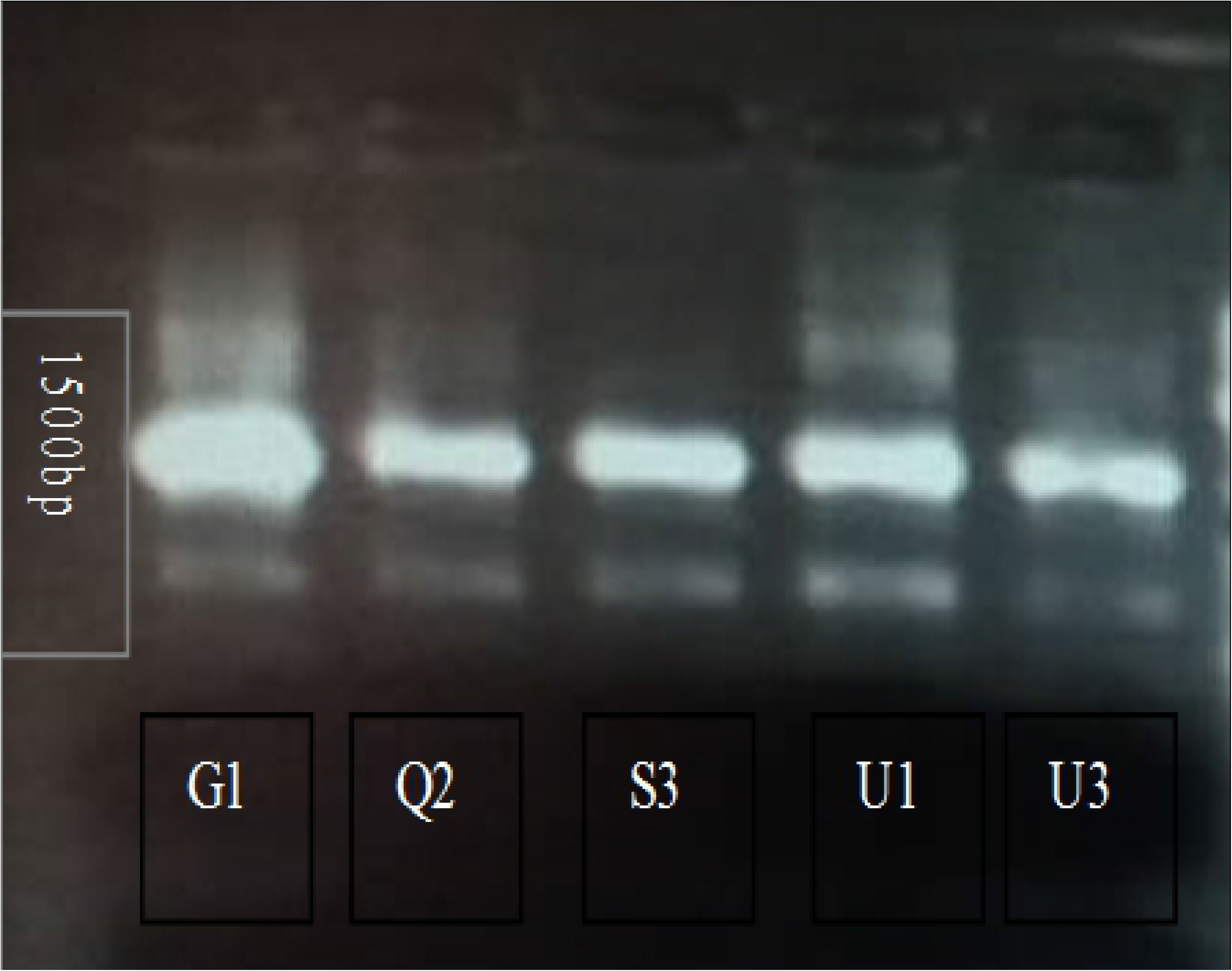
Gel electroporation of 16S rDNA amplified sequence.

### Phylogenetic analysis

The evolutionary history was inferred by using the Maximum Likelihood method based on the Tamura-Nei model. The percentage of trees in which the associated taxa clustered together is shown next to the branches. Initial tree(s) for the heuristic search were obtained automatically by applying Neighbor-Join and BioNJ algorithms to a matrix of pairwise distances estimated using the Maximum Composite Likelihood (MCL) approach, and then selecting the topology with superior log likelihood value. The tree is drawn to scale, with branch lengths measured in the number of substitutions per site. The analysis involved 25 nucleotide sequences. Codon positions included were 1st+2nd+3rd+Noncoding. There were a total of 1012 positions in the final dataset. Evolutionary analyses were conducted in MEGA X (Kumar *et al*., 2018, Fig 6).

**Fig 6.**
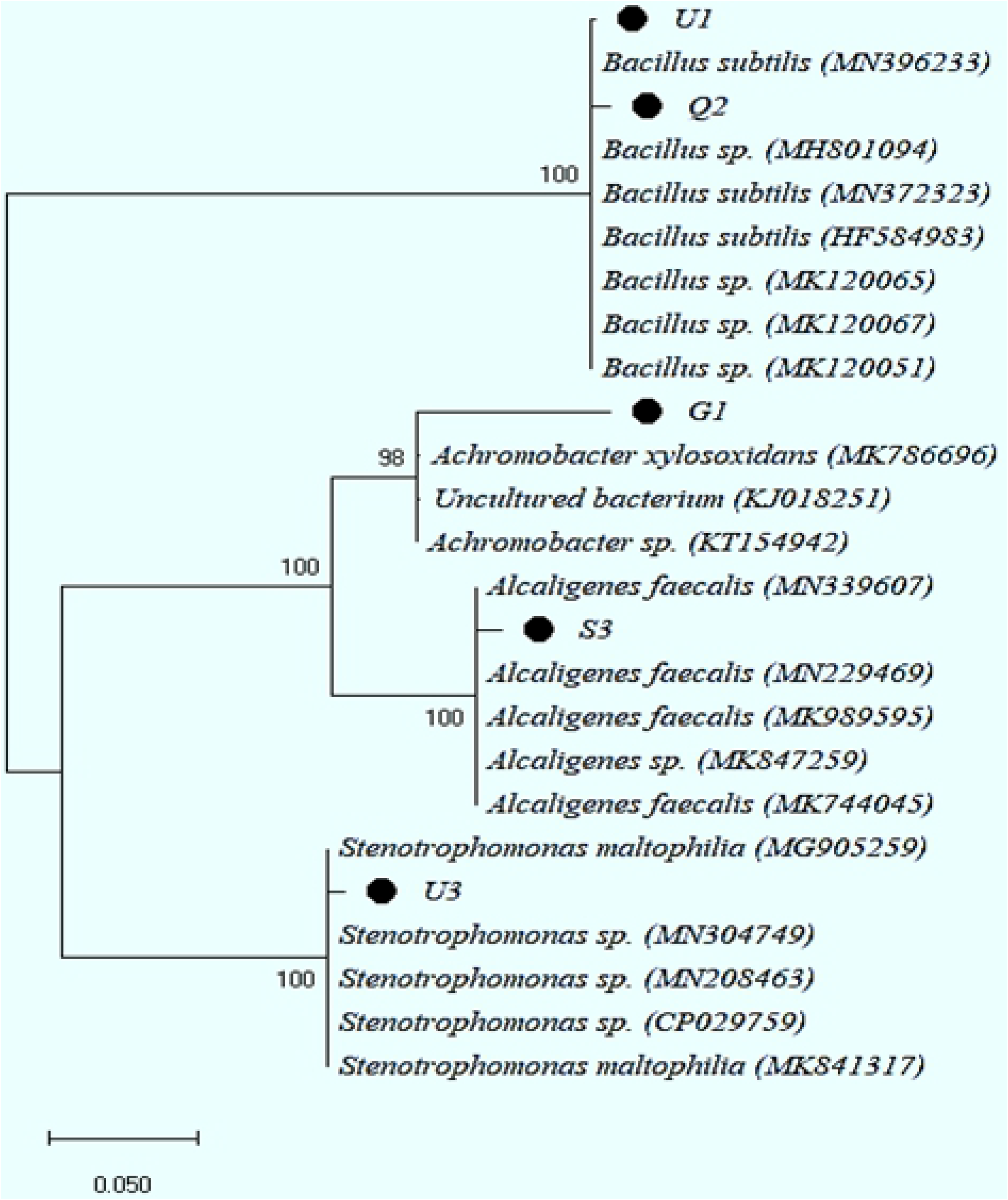
Phylogenetic tree for the bacterial isolates showing relatedness with other organisms.

### Microbial production of L-Glutaminase

Glutaminase production was done from all the bacterial isolates on Glutamine salt media and optimized for various production parameters.

### Optimization of parameters for glutaminase production

The optimization of glutaminase production by selected strain against different effectors like incubation time, effect of pH, effect of temperature carbon source, nitrogen source and inducers were carried out in shake flask experiments.

### Effect of incubation time

Glutaminase production was done from all the bacterial isolates on glutamine salt media at 30°C. Fig 7 illustrating amount of enzyme being produced from strains, which showed that *Achromobacter xylosoxidans* G1 produced highest amount of enzyme (44.13 IU/ml/min) at second day. At second day of incubation *Bacillus subtilis* Q2 was 37.30 IU/ml/min, *Stenotrophomonas maltophilia* U3 yield 41.92 IU/ml/min, *Bacillus subtilis* U1 35.3 IU/ml/min and *Alcaligenes faecalis* S3 exhibited high yield 39.71756399 IU/ml/min at 4^th^ day of incubation. At third and fourth day G1 and S3 produced high amount respectively.

**Fig 7.**
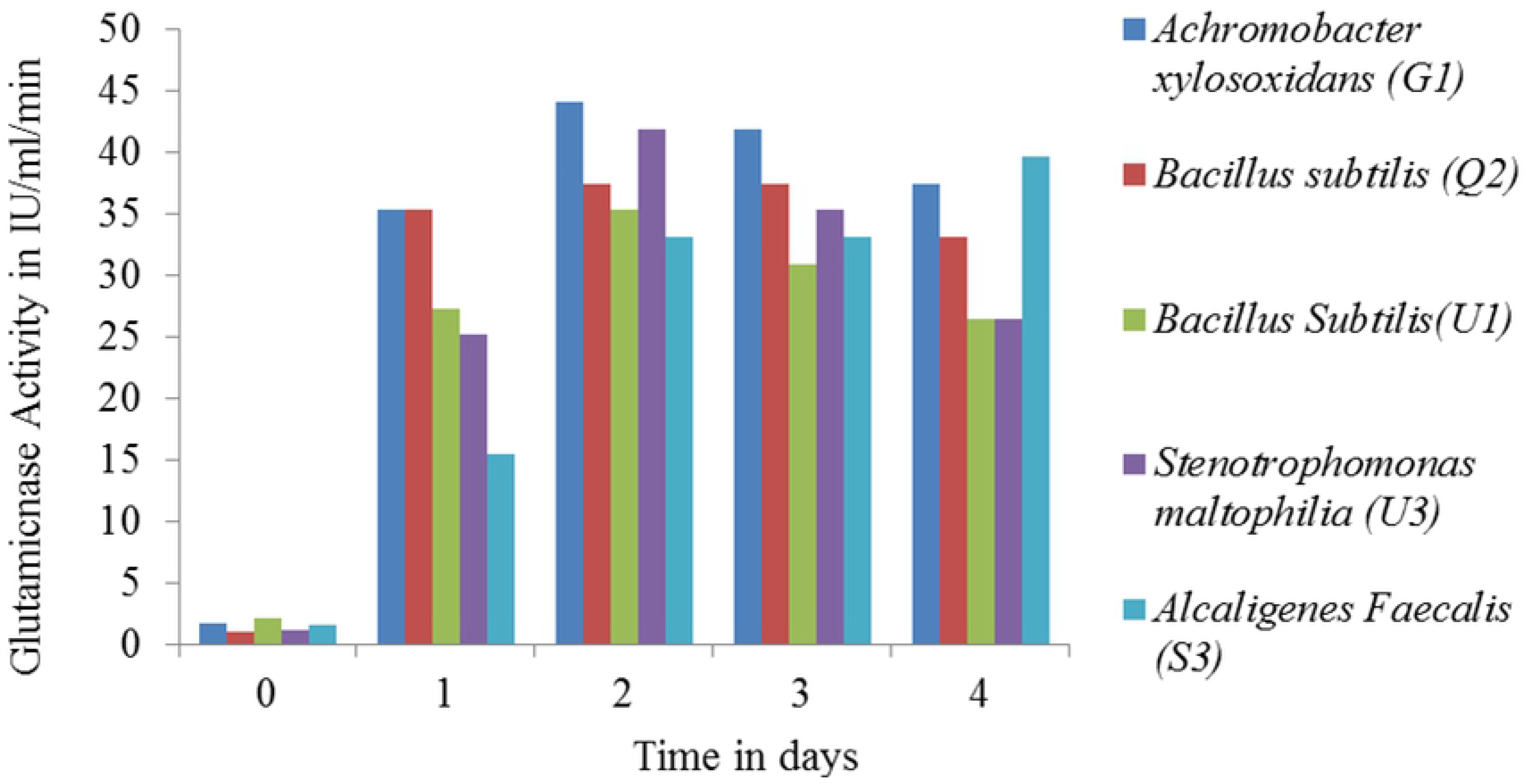
Production of glutaminases by different bacterial isolates in glutamine salt media incubated for 4 days.

### Effect of pH

Results showed that Glutaminase production by *Achromobacter xylosoxidans* G1, *Alcaligenes faecalis* S3 on production media at 37°C showed that optimum pH is 9. The strains *Bacillus subtilis* U1, showed maximum activity at pH 6, *Stenotrophomonas maltophilia* U3 shown best activity at pH 8 while *Bacillus subtilis* Q2 also showed glutamine activity between the pH range of 6-8 maximally (Fig 8). *Achromobacter xylosoxidans* G1 worked best giving activity of 57.369 U/ml/min at pH 9 at 3^rd^ day of incubation, the activity 29.126 U/ml/min of *Bacillus subtilis* Q2 was highest at pH 6 and 8 on 2^nd^ day, U3 showed maximum activity at pH 8 at third day that was 46.34 U/ml/min, *Bacillus subtilis* U1 showed highest activity of 37.069 U/ml/min at pH 6 while S3 showed highest activity of 57.36 U/ml/min at pH 9.

**Fig 8.**
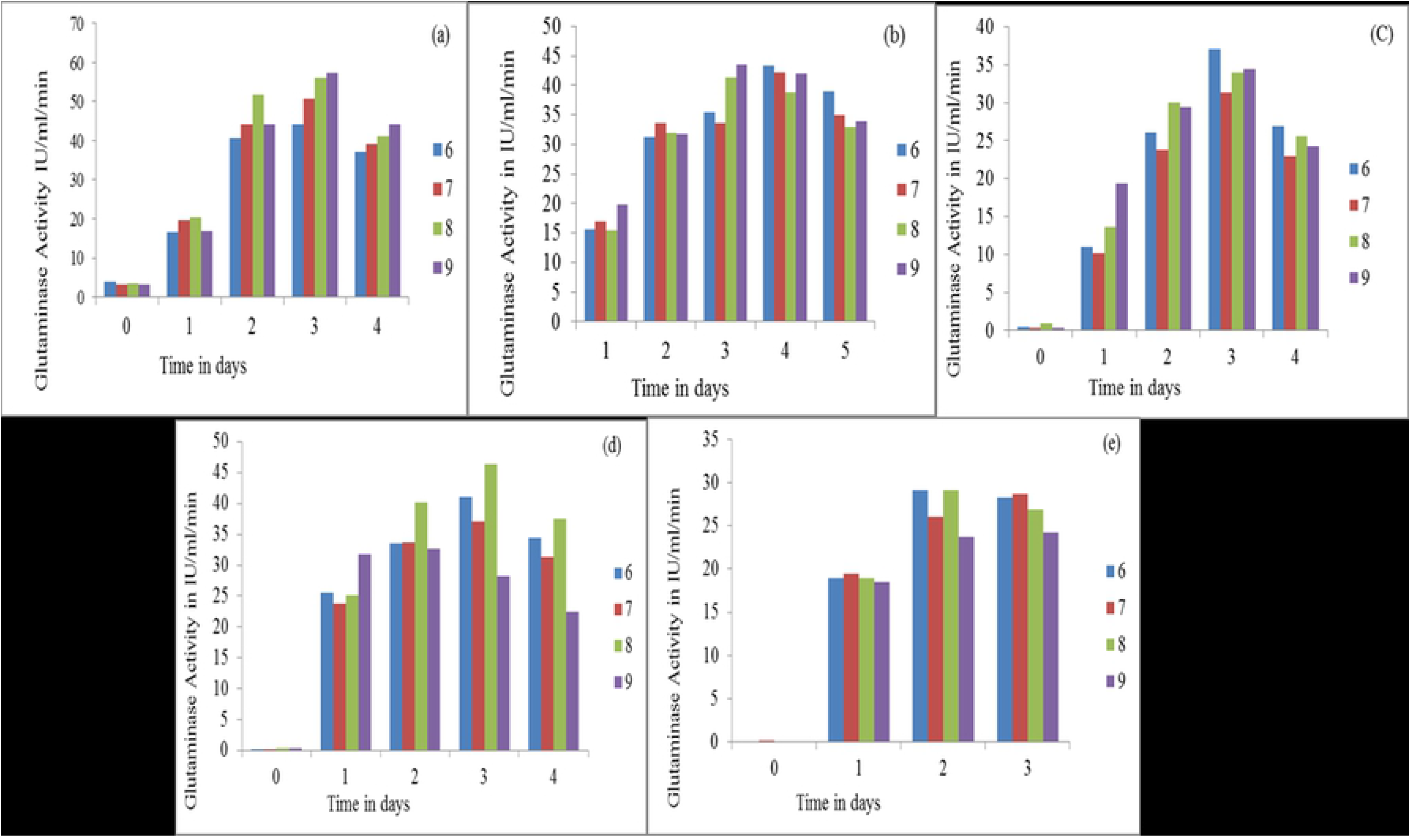
Effect of pH on glutaminase production by bacterial isolates; (a) *Achromobacter xylosoxidans* G1, (b) *Alcaligenes faecalis* S3, (c) *Bacillus subtilis* U1, (d) *Stenotrophomonas maltophilia* U3, (e) *Bacillus subtilis* Q2 on Glutamine Salt media.

### Effect of Temperature

Glutaminase production was observed at 25°C, 30°C and 37°C by all 5 selected bacterial strains. Highest Glutaminsae activity was shown at 37°C by *Bacillus subtilis* U1 achieved at 4^th^ day with 66.19 IU/ml/min and *Bacillus subtilis* Q2 (57.3 IU/ml/min) at 4^th^ day. The other strains *Achromobacter xylosoxidans* G1 and *Stenotrophomonas maltophilia* U3 showed maximum Glutaminase activity at 30°C at 3^rd^ day (44.1 IU/ml/min and 41.9 IU/ml/min respectively). While *Alcaligenes Faecalis* S3 best activity (39.71 IU/ml/min) was observed at 25°C at 3^rd^ day of incubation (Fig 9).

**Fig 9.**
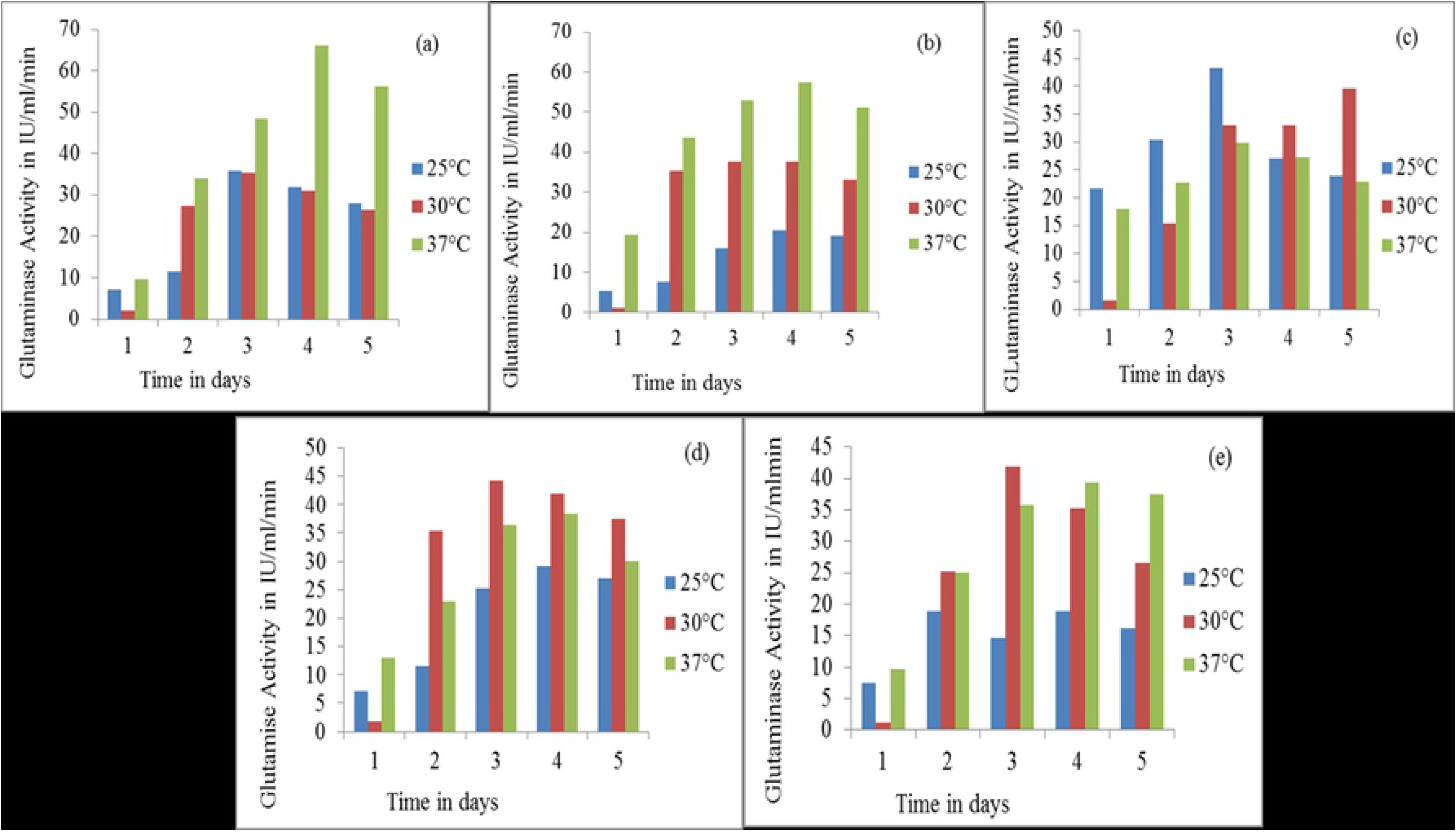
Effect of Temperature on Glutaminase production by bacterial isolates on Glutamine salt media. (a) *Bacillus subtilis* U1, (b) *Bacillus subtilis* Q2, (c) *Alcaligenes faecalis* S3, (d) *Achromobacter xylosoxidans* G1, (e) *Stenotrophomonas maltophilia* U3 on Glutamine Salt media.

### Effect of Carbon source

Enzyme from strain *Bacillus subtilis* U1 with carbon source showed high activity on 3rd and 4th day of incubation among which glucose was on highest influencing point of activity and sucrose was lowest activator. The strain *Stenotrophomonas maltophilia* U3 got highest activity with xylose as carbon source and showed least activity with glucose showing contrast feature with U1. The maximum activity achieved by *Bacillus subtilis* Q2 which showed a universal behavior regarding enzyme release under different carbon sources except Maltose. The highest in this case was also glucose. The best activators of enzyme were glucose (*Bacillus subtilis* U1 & *Bacillus subtilis* Q2), sorbitol (*Alcaligenes faecalis* S3 & *Achromobacter xylosoxidans* G1) and xylose (*Stenotrophomonas maltophilia* U3) showing activities of 50.8, 60.9, 42.674, 29 and 47.66 IU/ml/min respectively (Fig 10).

**Fig 10.**
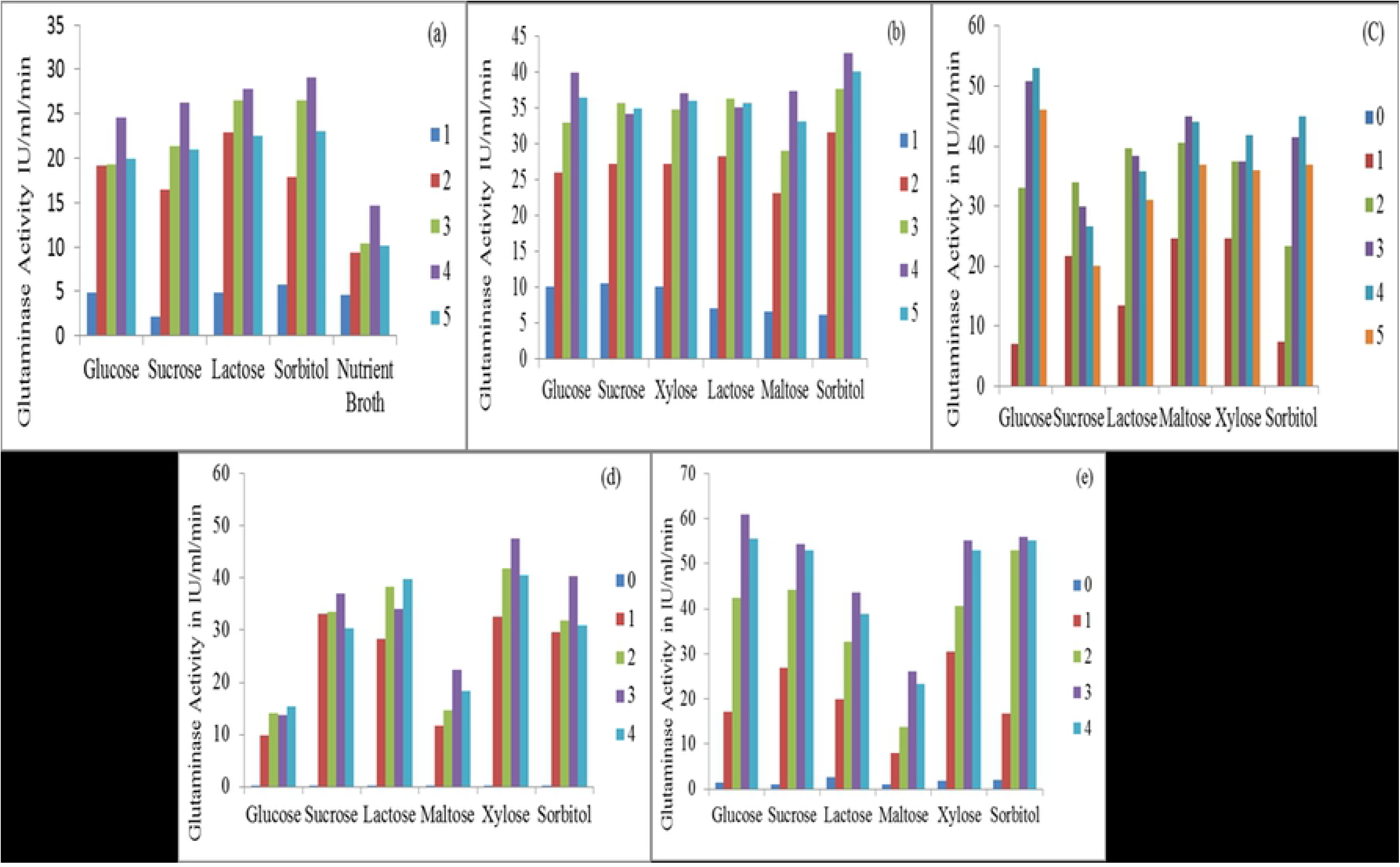
Effect of Carbon source (200mM 5ml/50ml) on Glutaminase production by bacterial isolates in fermentation media at 37°C; a strain (a) *Achromobacter xylosoxidans*

### Effect of Nitrogen source

The production of L-glutaminase by strain *Achromobacter xylosoxidans* G1 was at its higher peak when yeast extract was used as nitrogen source showing 22.947 IU/ml/min activity (Figure 11). The poorest nitrogen source was sodium nitrate and ammonium sulphate at day 1 and day 2 respectively. In the case of *Bacillus subtilis* U1 the Trypton was the best nitrogen donor showed maximum activity up to 61.7828 U/ml/min (Fig 11).

**Fig 11.**
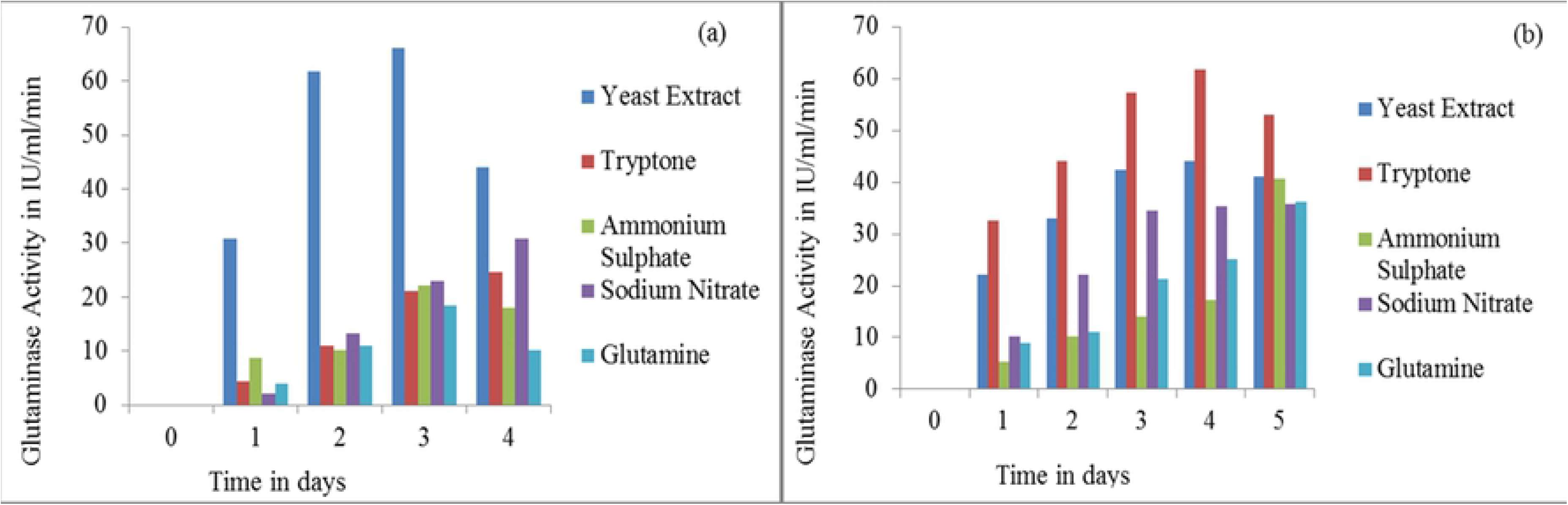
Effect of Nitrogen source (200mM 4ml/50ml) for Glutaminase production by G1 (a) *Archomobacter xylosoxidans* G1, (b) *Bacillus subtilis* U1 on Mineral salt Media.

### Effect of Inducers

The effect of inducers on all the strains were observed for 4 days. The peak achieved at 3^rd^ day with strain U1 (Fig 12).

**Fig 12.**
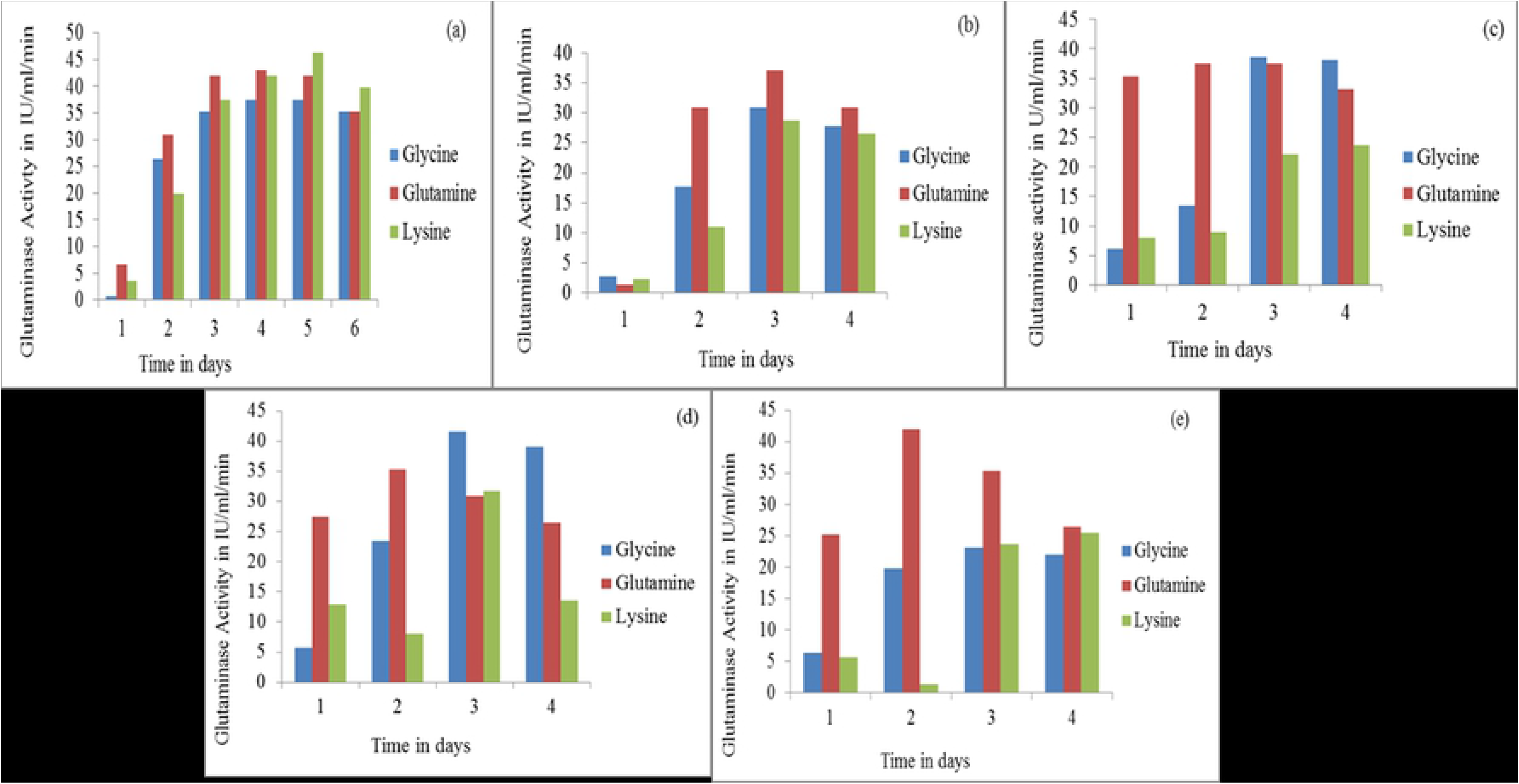
Effect of Inducers on Glutaminase production by bacterial isolates in (a) *Achromobacter xylosoxidans* G1, (b) *Alcaligenes faecalis* S3, (c) *Bacillus subtilis* Q2, (d) *Bacillus subtilis* U1, (e) *Stenotrophomonas maltophilia* U3 on Glutamine salt media

The inducers such as Glycine, Glutamine and Lysine influenced the glutaminase activity in a strain G1. The interesting feature was that all the inducer showed parabolic effect on enzyme’s activity such that increasing at starting point and showed highest activity at 3rd to 5th day and ultimately decreased at 6th day. The glutamine tends to effect greatly on all of strain like its activities in U3, S3 and Q2 were 41.92, 37.069 and 38.6 U/ml/min at day 2^nd^, 4^th^ and 3^rd^ day except G1 where lysine enhanced the activity by 46.33 IU/ml/min of enzyme and *Bacillus subtilis* U1 showed maximum activity with glycine (41.48 IU/ml/min) (Fig 12).

## Discussion

Since, the disclosure of L-glutaminase for its properties like anti cancerous, different microbial sources is the focal point of enthusiasm for the isolation of the protein. L-glutaminase action is studied in living organisms, plant tissues and microorganisms including microscopic organisms, and actinomycetes. Microbial L-glutaminase (L-glutamine amido hydrolase EC 3.5.1.2) has gotten more prominent consideration for its potential biotechnological applications and effectiveness in wide range.

In the present study we have isolated 4 bacterial strains *Bacillus subtilis* U1, *Stenotrophomonas maltophilia* U3, *Bacillus subtilis* Q2 from different soil samples and *Achromobater xylosoxidane* G1 that was isolated from the old L-Glutamine sample. In general, glutaminases from *E. coli*, *Bacillus spp* [20] *Pseudomonas spp*., *Citrobacter*, *staphylococcus* [21] have been isolated and well-studied.

The primary screening was performed on Glutamine salt media total 20 strains were isolated initially out of which we have selected five isolates giving good activities in screening assays giving pink color in the medium containing phenol red as indicator. Investigations of Aly [22] indicated the amidase presence and activity in many organisms. Emelda [23] isolated bacteria from Soil and aquatic environment screened on minimal glutamine media with phenol red as an indicator and selected the colonies produced pink color due to release of ammonia into the media.

In the bacterial screening with substrate all of the selected strains showed positive results *Bacillus subtilis* U1 showed largest zone 34 mm while *Bacillus subtilis* Q2 gave smallest zone of (16 mm) in screening media while testing the zone of hydrolysis with cell free supernatants of the isolates. This was trailed by the isolation of glutaminases from different microbial sources for restorative applications [24, 25].

Taxonomic characterization of all the selected isolates were performed and all the biochemical tests were positive except glucose fermentation in G1, MSA and indole tests in *Alcaligenes faecalis* S3 and Urease in *Bacillus subtilis* Q2, catalase, citrate urease and indole in *Bacillus subtilis* U1. The strain *Stenotrophomonas maltophilia* U3 was negative for all of biochemical tests. L-Glutaminase was isolated from *Bacillus subtilis* which gave 50% tests positive when 34 biochemical tests were performed [25].

The sequencing of L-Glutamisae producing strains of U1 showed 98.32% homology with *Bacillus subtilis* strain B92, S3 has shown 97.87% homology with *Alcaligenes faecalis* strain SDU20, G1 was 92.35% homologous to *Achromobacter xylosoxidans* strain BPS6, Q2 was 97.87% was homologous to *Bacillus subtilis* strain B18 and U3 was 96.56% was homologous to the *Stenotrophomonas maltophilia* strain NA156 on the basis of 16S ribosomal RNA gene partial sequence. Molecular identification of 16S rDNA analysis of L-glutaminase producing bacteria identified as *Bacillus subtilis* JK-79 [26]. *Alcaligenes faecalis* KLU102 was isolated from marine for glutaminase production [27]. *Stenotrophomonas maltophilia* and *Achromobacter* specie have been reported for L-glutamiase production [28] [29].

Selected bacterial strains were optimized for incubation time, pH, temperature, carbon source, nitrogen source and effects of inducers. For incubation time *Achromobacter xylosoxidans* G1, *Bacillus subtilis* Q2, *Stenotrophomonas maltophilia* U3 and *Bacillus subtilis* U1 produced highest glutaminase at second day while *Alcaligenes faecalis* S3 showed highest activity 39.7 IU/ml/min at 4^th^ day of incubation. Maximum L-glutaminase production was achieved in submerged fermentation after 18 hours of incubation time by marine isolated *Bacillus subtilis* [30] and 72 hours of incubation for *Pseudomonase VJ-6* [31].

The L-Glutaminase activity by *Achromobacter xylosoxidans* G1, *Bacillus subtilis* Q2, *Stenotrophomonas maltophilia* U3, *Bacillus subtilis* U1 and *Alcaligenes faecalis* S3 was noted to be best at pH 9, 6, 8, 6, 9 with respectively. Best L-glutaminase production was observed between pH 7 from the forest soil isolated bacterial strain of *Bacillus* sp [32]. Marine isolated *Vibrio azureus* JK-79 shown maximum glutaminase production at pH 8 [33].

Bacterial strains were optimized for glutaminase production at 25°C, 30°C and 37°C. Glutaminsae activity was maximum at 37°C by *Bacillus subtilis* U1 (66.19 IU/ml/min) and *Bacillus subtilis* (Q2) 57.3 IU/ml/min and other strains *Achromobacter xylosoxidans* (G1) and *Stenotrophomonas maltophilia* (U3) showed best activities at 30C° (Figure 9). While for *Alcaligenes Faecalis* (S3) 25°C was the optimum temperature for production. Kiruthika [33] isolated *Vibrio azureus* JK-79 from marine, shown maximum glutaminase activity at 37°C. Al-Zahrani [34] found 35°C is the best temperature for glutaminase production by *Psuedomonas* NS16.

Glucose for *Bacillus subtilis* U1 and *Bacillus subtilis* Q2, sorbitol for *Alcaligenes faecalis* S3 and *Achromobacter xylosoxidans* G1, xylose for *Stenotrophomonas maltophilia* U3 were the best carbon source for glutaminase production when different carbon sources were tested (Figure 10). The present study focuses on the selective isolation of the potent L-glutaminase producing soil bacteria. Glucose was best carbon source for glutaminase production by *Pseudomonase aurignosa* [34].

The best nitrogen source was yeast extract G1 and Trypton for U1. Maximum glutaminase activity by *Pseudomonas aurignosa* was achieved with glutamine out of various nitrogen sources [34]. Kiruthika [35] identified glutaminase production by marine *Bacillus subtilis* JK-79 was enhanced by yeast extract.

The best inducer for *Bacillus subtilis* U1 was glycine while glutaminase production by *Stenotrophomonas maltophilia* U3 was equally induced by glycine and glutamine, *Alcaligenes faecalis* S3 and *Bacillus subtilis* Q2 showed best activities with glutamine while for *Achromobacter xylosoxidans* G1 it was lysine. Glutaminase production was induced by glutamine in case of *Bacillus subtilis* JK-79 and *Bacillus* sp. [35] [36].

## Conclusion

Microbial L-glutaminases with improved properties like universal presence, thermo-resistance and salt resilience discovers applications in nourishment industry just as in malignant growth treatment. Utilization of atomic instruments like site coordinated mutagenesis, coordinated advancement for the improvement of L-glutaminase is still at outset and further engaged research is required toward this path. Wide extension still exists on the screening and choice of wild and novel glutaminase creating life forms from different environmental specialties for the generation of L-glutaminase and their further improvement with the assistance of present-day advances [37]. Glutamine is an incredibly adaptable supplement that adds to numerous parts of mediator digestion in malignant growth cells. It is especially significant in the development of the macromolecules required for cell multiplication and protection from oxidative pressure. Since glutamine digestion is modified during dangerous change, imaging techniques that target glutamine should give a helpful window into tumor science that would supplement.

There is need of combination or elaboration of such system or pathways which turn away or make malignant growth cell dead as well as purpose recovery impact for typical body cells and improve conventional execution or working of standard cell by advancing their immunology at sub-atomic level [38].

